# Preserving protein homeostasis prevents motor impairment in DNA Damage Response-compromised *C. elegans*

**DOI:** 10.1101/2021.12.22.473820

**Authors:** Wouter Huiting, Alejandra Duque-Jaramillo, Renée I. Seinstra, Harm. H. Kampinga, Ellen A.A. Nollen, Steven Bergink

## Abstract

To maintain genome integrity, cells rely on a complex system of DNA repair pathways and cell cycle checkpoints, together referred to as the DNA damage response (DDR). Impairments in DDR pathways are linked to cancer, but also to a wide range of degenerative processes, frequently including progressive neuropathy and accelerated aging. How defects in mechanistically distinct DDR pathways can drive similar degenerative phenotypes is not understood. Here we show that defects in various DDR components are linked to a loss of protein homeostasis in *Caenorhabditis elegans*. Prolonged silencing of *atm-1, brc-1* or *ung-1*, central components in respectively checkpoint signaling, double strand break repair and base excision repair enhances the global aggregation of proteins occurring in adult animals, and accelerates polyglutamine protein aggregation in a model for neurodegenerative diseases. Overexpression of the molecular chaperone HSP-16.2 prevents enhanced protein aggregation in *atm-1, brc-1* or *ung-1*-compromised animals. Strikingly, rebalancing protein homeostasis with HSP-16.2 almost completely rescues age-associated impaired motor function in these animals as well. This reveals that the consequences of a loss of *atm-1, brc-1* or *ung-1* converge on an impaired protein homeostasis to cause degeneration. These findings indicate that a loss of protein homeostasis is a crucial downstream consequence of DNA repair defects, and thereby provide an attractive novel framework for understanding the broad link between DDR defects and degenerative processes.

## INTRODUCTION

To maintain genome integrity, cells rely on the various DNA repair pathways and cell cycle checkpoints together referred to as the DNA damage response (DDR) (Jackson & Bartek, 2009). Failure or dysregulation of DDR processes can have dramatic consequences, which is underlined by the more than 50 human disorders caused by inherited DDR defects (Petr et al., 2020). These disorders, linked to mutations in genes that function in mechanistically distinct DDR pathways, are often characterized by a range of different pathological features, including cancer predisposition, neurodegeneration, and accelerated ageing. This has fueled the idea that a decline of genome maintenance is a driving force of the degenerative processes that occur with ageing (Schumacher et al., 2021). However, how DDR defects can drive a range of disease phenotypes and tissue degeneration besides cancer, particularly in the early stages preceding cell death, is still largely unclear.

Greater insight may be provided by a primary feature of ageing tissues, which is a gradual loss of protein homeostasis (Labbadia & Morimoto, 2015). Over time, the human proteome tends to drift from a state of balance towards a more instable state. This loss of protein homeostasis is hallmarked by a global increase in dysregulated and misfolded proteins that, if not properly degraded, can self-assemble into potentially cytotoxic protein aggregates (reviewed in (Hipp et al., 2019)). A loss of protein homeostasis is associated with cellular dysfunction, and thought to be causative for a range of – often degenerative – disorders (i.e. ‘proteinopathies’), including several (cardio)myopathies, kidney diseases, and most notably neurodegenerative disorders like Alzheimer’s (AD), Parkinson’s (PD), and Huntington’s diseases (HD) (reviewed in (Klaips et al., 2018; Labbadia & Morimoto, 2015)). Accumulating evidence also links impairments in DDR pathways to proteinopathies (Bettencourt et al., 2016; Huiting & Bergink, 2021; Sepe et al., 2016; Shackelford, 2006; Suberbielle et al., 2015; Weissman et al., 2007; Yu et al., 2018). While most of this evidence is correlative in nature, defects in the checkpoint kinase ATM (ataxia telangiectasia mutated) have recently been shown to result in a loss of protein homeostasis. Based on these findings we hypothesize that there is a causal relationship: defects in certain DDR pathways challenge protein homeostasis, and this contributes to the occurrence of degenerative phenotypes.

Here, we report that disruption of various DDR pathways exacerbates the age-dependent loss of protein homeostasis in *Caenorhabditis elegans*. Remarkably, mitigating protein aggregation is sufficient to prevent the motor impairment that we observed in animals suffering from DDR defects. Thus, this suggests that the loss of protein homeostasis that we find upon impairments in several DDR pathways is an important downstream effector that drives degeneration.

## RESULTS

### Silencing of distinct DDR components exacerbates endogenous protein aggregation

To investigate the relationship between defects in DDR pathways and protein homeostasis, we turned to the invertebrate model organism *C. elegans*, a nematode that shares many evolutionary conserved pathways with humans, including those involved in the DDR (Lans & Vermeulen, 2015). It has a largely post-mitotic soma, which makes it an ideal model to assess long-term consequences of DDR defects on the proteome. *C. elegans* displays a loss of protein homeostasis during ageing, hallmarked by a global accumulation of insoluble, aggregated proteins (David et al., 2010; Walther et al., 2015). Protein aggregation can be used as a proxy to investigate the state of protein homeostasis in animals (Alavez, 2017), and aggregated proteins can be isolated through detergent fractionation (David et al., 2010; Groh et al., 2017; Huang et al., 2019; Walther et al., 2015). In our study, we used a 1% SDS fractionation method to extract an insoluble protein fraction from animals (Supplemental Figure 1A, B), which we from here on refer to as aggregated proteins. We first tested if knockdown of the checkpoint signaling kinase ATM-1 (the ATM homologue in *C. elegans*) increased protein aggregation, as it does in human and yeast (Corcoles-Saez et al., 2018; Huiting et al., 2021; Lee et al., 2018). A feeding RNAi method in Bristol N2 wild-type animals was used to induce a knockdown of ATM-1 from L1 larval stage onwards (Figure 1A, Supplemental Figure 1D). Knockdown of ATM-1 resulted in a significant increase in protein aggregation as compared to animals fed the empty vector (EV ctrl) (Figure 1B-D). The increased aggregation after knockdown of ATM-1 seems to be due to the loss in DDR-functioning as knockdown of RAD-50 and MRE-11, which function upstream of ATM-1 in the DDR, also led to an increase in aggregated proteins (Figure 1B, Supplemental Figure 1C), as has recently been found for MRE-11 in human cells as well (Lee et al., 2021).

**Figure 1.**
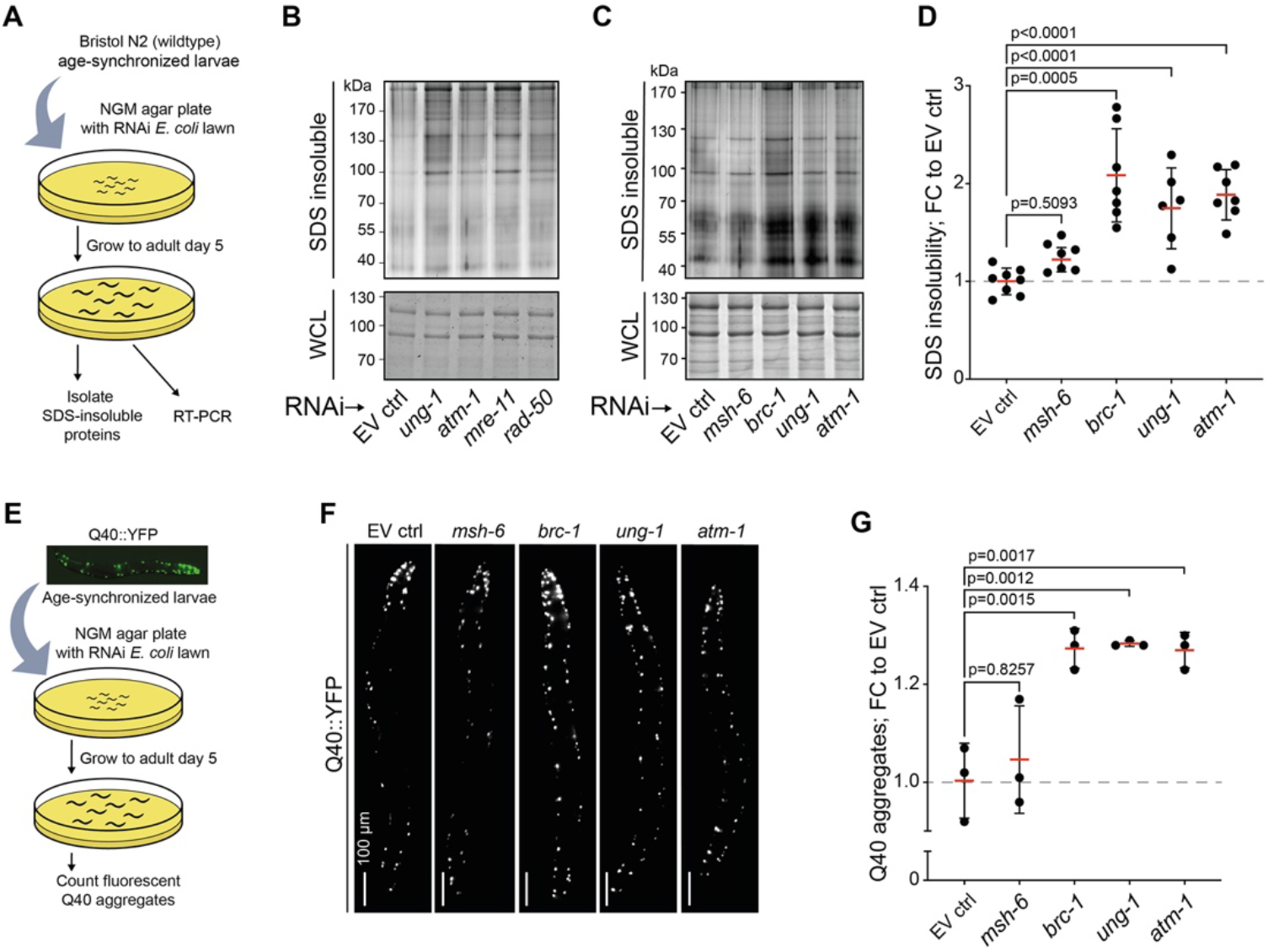
Protein aggregation is increased in DDR-compromised *C. elegans*. (A) Experimental outline. (B-C) SDS-insoluble and WCL protein fractions of animals fed RNAi’s targeting the indicated DDR components, run on SDS-PAGE gels and subjected to in-gel silver or Coomassie staining, respectively. (D) Densitometric quantification of C. Values represent the fold change (FC) of the amount of SDS-insoluble normalized to the empty vector control. Bars represent mean ± SD (standard deviation). Significance calculated using a one-way ANOVA with Dunnet’s post-hoc test. (E) Experimental outline. (F) Typical fluorescence stereomicroscopy pictures of AM141 (rmIs133 [unc-54p::Q40::YFP]) animals subjected to EV ctrl or *msh-6, brc-1, ung-1* or *atm-1* RNAi. (G) Quantification of Q40::YFP aggregates, relative to EV ctrl animals. Bars represent mean ± SD. Significance calculated using a one-way ANOVA with Dunnet’s post-hoc test. In all graphs, circles represent independent biological repeats.

We also tested if silencing other, distinct DDR genes led to an increase in insoluble proteins (see also Supplemental Table 1 for an overview of all DDR genes targeted in this study): MutS homologue in mismatch repair *msh-6*, the base-excision repair glycosylase *ung-1*, and the double strand break repair E3 ligase *brc-1* (Supplemental Figure 1D). Interestingly, gene knockdown of *ung-1* and *brc-1* exacerbated protein aggregation as well (Figure 1B-D). Knockdown of *msh-6* had no significant effect on protein aggregation (Figure 1C, D).

Various proteinopathies, including Huntington’s disease (HD) and several spinocerebellar ataxias (SCAs), are primarily caused by proteins containing expanded polyglutamine stretches (polyQ) that are aggregation-prone (Kuiper et al., 2017; Lieberman et al., 2019). As these polyQ-proteins are known to aggregate faster in the context of a disrupted protein homeostasis (Gidalevitz, 2006; Nollen et al., 2004), their aggregation can be used as another protein homeostasis read-out. We subjected animals expressing a Q40::YFP construct (from here on referred to as polyQ) in their body wall muscle cells (strain AM141 (Morley et al., 2002)) to the same RNAi-mediated knockdown regime as before (Figure 1E). At adult day 5, polyQ aggregates were scored by counting the number of fluorescent foci per animal (Figure 1F). Knockdown of the DDR genes *brc-1, atm-1* and *ung-1* resulted in a significant ∼25% increase in the number of polyQ aggregates (Figure 1F, G; Supplemental Figure 1E). In contrast, targeting *msh-6* did not result in an increased number of polyQ foci. Knockdown of none of the DDR genes significantly altered polyQ protein levels compared to EV ctrl animals (Supplemental Figure 1F, G), indicating that the enhanced polyQ aggregation is not caused by polyQ expression changes. These findings support the protein insolubility data, together suggesting that impairments in several different DDR pathways over time result in a disrupted protein homeostasis in *C. elegans*.

### The small heat shock protein HSP-16.2 maintains protein homeostasis upon an impaired

*DDR* We previously found that a loss of protein homeostasis in ATM-deficient mammalian cells is associated with an increased expression of the small heat shock protein HSPB5 (also known as αB-crystallin), and that overexpression of HSPB5 is able to suppress aggregation caused by ATM-deficiency (Huiting et al., 2021). The *C. elegans* HSP-16 protein family is considered to be functionally orthologous to human HSPB5 (Ganea, 2001). The best studied member of this family is HSP-16.2, whose expression levels have been shown to correlate with worm life span (Rea et al., 2005). Moreover, overexpression of HSP-16.2 in *C. elegans* suppresses toxic protein aggregation and promotes longevity (Fonte et al., 2008; Walker & Lithgow, 2003). To test whether overexpression of HSP-16.2 can also mitigate the accelerated protein aggregation caused by a compromised DDR, we employed a strain harboring an extra GFP-tagged copy of HSP-16.2 under the control of its endogenous promoter. This strain (GL400) has been shown to be long-lived and suffer less from proteotoxic stress (McColl et al., 2010). Using the same RNAi feeding strategy as before we targeted *atm-1, brc-1, ung-1* and *msh-6*, in GL400 animals and in a matched wild-type strain HE1006 (Supplemental Figure 2A). Interestingly, knockdown of *ung-1, brc-1* as well as *atm-1* in GL400 animals resulted in an increased expression of HSP-16.2 compared to EV ctrl animals, primarily in the intestinal region (Figure 2A). This is similar to what can be observed upon treating GL400 animals with a proteotoxic heat shock treatment (Supplemental Figure 2B) that also disturbs protein homeostasis and is associated with exacerbated protein aggregation (Zevian & Yanowitz, 2014). In contrast, no increase in HSP-16.2 signal was seen upon knockdown of *msh-6* (Figure 2A). We quantified protein aggregation from HE1006 wild-type and GL400 HSP-16.2 post-replicative animals. The HSP-16.2 overexpression strain was resistant to the increased protein aggregation caused by knockdown of *ung-1, atm-1* or *brc-1* (Figure 2B, C), despite comparable knockdown efficiencies for each gene between the two strains (Supplemental Figure 2A). From this, we conclude that HSP-16.2 is able to buffer the exacerbated protein aggregation occurring in animals subjected to a knockdown of *ung-1, atm-1* or *brc-1*.

**Figure 2.**
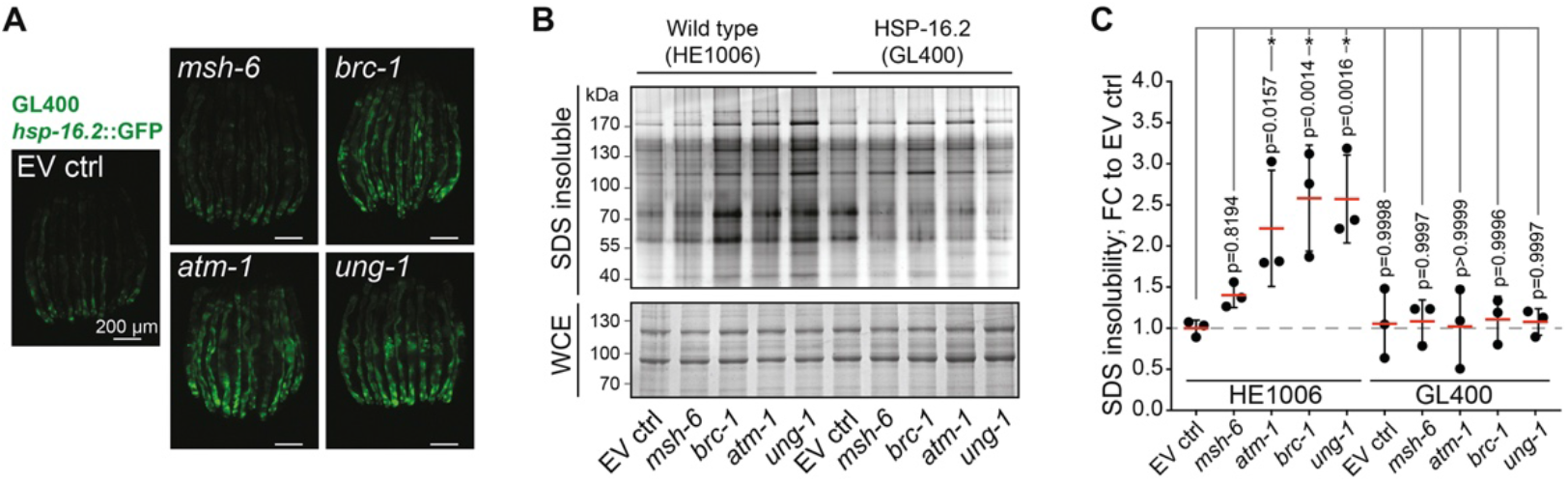
HSP-16.2 is able to guard protein homeostasis in DDR-compromised *C. elegans*. (A) HSP-16.2::GFP signal in (adult day 3) GL400 animals subjected to EV ctrl or *msh-6, atm-1*-, *ung-1* or *brc-1* RNAi. Typical fluorescence stereomicroscopy pictures are shown. (B) SDS-insoluble and WCL fractions of EV ctrl animals and animals fed *msh-6, brc-1, atm-1* and *ung-1* RNAi in HE1006 (wild-type) and GL400 (HSP-16.2 overexpression) animals. (C) Densitometric quantification of B. Values represent the fold change (FC) of the amount of SDS-insoluble normalized to the EV control. Bars represent mean ± SD. Significance calculated using a one-way ANOVA with Dunnet’s post-hoc test.

To our knowledge neither HSPB5/αB-crystallin nor any of its orthologs – including the *C. elegans* HSP-16 family – have been reported to play a role in DNA repair or checkpoint signaling (Huiting et al., 2021). We nevertheless investigated whether the decreased protein aggregation might be caused by an unanticipated direct impact of HSP-16.2 on the DDR. We first tested whether the presence of an extra copy of *hsp-16.2* affects gamma irradiation-induced cell death in the germline, an established early read-out for impaired genome maintenance in *C. elegans* (Craig et al., 2012). Using acridine orange, a dye that is selectively retained in apoptotic cells (Lant & Derry, 2014), no differences in cell death between untreated HE1006 wild-type and GL400 HSP-16.2 animals for any of the RNAi’s tested were found (Supplemental Figure 2C, D). In all conditions irradiation with 100 gray resulted in an increase in acridine-positive cells, reflecting the increase in germline apoptosis after DNA damage (Supplemental Figure 2C). In line with a DNA repair defect, knockdown of *ung-1* and *brc-1* resulted in a higher sensitivity towards DNA damage, as indicated by the relative increase in acridine-positive cells after gamma irradiation (Supplemental Figure 2E). *Atm-1* knockdown animals did not display this increase, reflecting the critical role of ATM in DNA damage-induced apoptosis (Shiloh & Ziv, 2013)(Supplemental Figure 2E). Importantly, HSP-16.2 animals did not show an altered response after γ-irradiation, under any RNAi condition tested (Supplemental Figure 2C, D). Certain DNA repair defects are also known to cause a developmental delay in *C. elegans* as a consequence of an accumulation of DNA damage (Lans & Vermeulen, 2015). We noted a clear developmental delay in age-synchronized offspring of HE1006 wild-type animals subjected to *brc-1* RNAi in particular. Whilst HSP16.2 reduced the protein aggregation induced by *brc-1* RNAi in adult postmitotic worms, no difference was found in the extent of developmental growth delay between wild-type animals and GL400 HSP-16.2 animals subjected to *brc-1* RNAi (Supplemental Figure 2F). Together, these findings argue that HSP-16.2 does not affect the capacity of DDR processes itself, but rather acts at a proteome level to buffer the exacerbated protein aggregation caused by a long-term disruption of various DDR pathways.

### Guarding protein homeostasis with HSP-16.2 is sufficient to prevent motor impairment in DDR-compromised animals

In several model organisms, including *C. elegans*, an age-related disruption of protein homeostasis is an important driver of the gradual decline in muscle function and movement (Ben-Zvi et al., 2009; Demontis & Perrimon, 2010; Huang et al., 2019). We investigated whether the enhanced protein aggregation caused by the silencing of DDR components accelerates this age-related motor impairment, and if elevated HSP-16.2 protects against such effects. We first evaluated if depletion of *atm-1, brc-1* or *ung-1* induces motor impairment by measuring crawling activity on adult day 5, the same timepoint at which we found that protein aggregation is enhanced (Figure 3A). In HE1006 wild-type animals, crawling activity was 20-30% lower after a knockdown of *atm-1, brc-1* and *ung-1* compared to EV ctrl (Figure 3B, C). GL400 HSP-16.2 outperformed the HE1006 wild-type animals, in line with the increased health span that has been reported before (McColl et al., 2010). However, normalization reveals that the presence of an extra *hsp-16.2* copy in the GL400 strain significantly reduced this relative motor defect in *atm-1* and *brc-1* knockdown animals, and restored crawling activity back to EV ctrl levels (Fig 3B, C).

**Figure 3.**
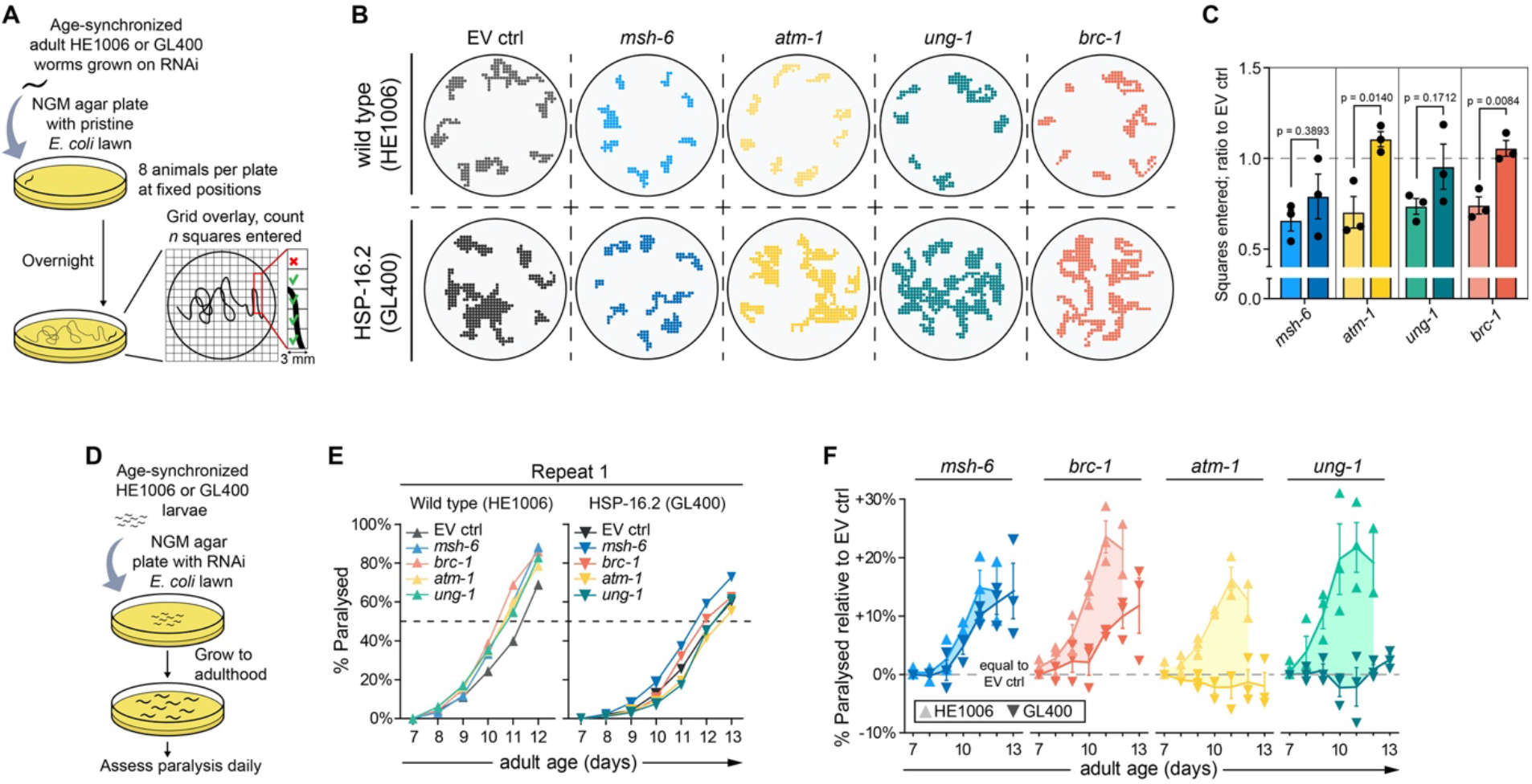
HSP-16.2 prevents a motor impairment in DDR-compromised *C. elegans*. (A) Experimental outline. (B) Representative result of individual trajectories of HE1006 (wild-type) and GL400 (HSP-16.2 overexpression) animals subjected to EV ctrl or the indicated RNAi’s. Tracks of 8 animals per plate are shown, of one representative experiment. (C) Quantification of B compared to each respective EV ctrl. Significance calculated using Student’s t-tests. Data is represented as mean ± SEM (standard error of mean). Circles represent independent biological repeats. (D) Experimental outline. (E) Paralysis rates of HE1006 wild-type (left) and GL400 HSP-16.2 overexpression (right) animals fed the indicated RNAi bacteria. One representative experiment is shown. (F) Difference in the percentage of paralyzed animals compared to EV ctrl over time, in both HE1006 (wild-type) and GL400 (HSP-16.2 overexpression) strains. >0% indicates a faster paralysis rate than EV ctrl animals. Triangles represent the mean result of each independent biological repeat. Data is represented as mean ± SEM.

We next used paralysis as a proxy for motor impairment over time (Figure 3D). Knockdown of *brc-1, atm-1* and *ung-1* in the wild type strain accelerated relative paralysis rates with age. In contrast, HSP-16.2 overexpression substantially or even completely protected against the accelerated paralysis induced by the silencing of these three genes (Figure 3E, F).

Interestingly, in both crawling and paralysis assays knockdown of *msh-6* also resulted in a lower motor performance, but in neither of these overexpression of HSP-16.2 was protective (Figure 3B, C and E, F). As we previously found that a knockdown of *msh-6* had no significant effect on protein aggregation (Figure 1D, G; Figure 2C) or on the fluorescent HSP-16.2 signal (Figure 2A), this indicates that targeting *msh-6* from the L1 larval stage onwards causes a motor defect largely unrelated to protein homeostasis. These findings are important, because they show that HSP-16.2 is able to rescue motor impairment induced by defects in DDR pathways specifically and only when those defects lead to a disruption of protein homeostasis.

## DISCUSSION

Our findings reveal that silencing of *atm-1, brc-1* or *ung-1* results in a motor impairment in *C. elegans* that is associated with a disruption of protein homeostasis. This loss of protein homeostasis is hallmarked by an increase in global protein aggregation, enhanced polyQ puncta formation, and up-regulated HSP-16.2 expression. The finding that HSP-16.2 overexpression reduces both the enhanced aggregation and restores motor impairment reveals a causality between the two. This points for the first time at the possibility that long-term consequences of defects in at least certain DDR pathways cause age-related degeneration by affecting protein homeostasis.

ATM-1, BRC-1 and UNG-1 fulfill largely different roles in distinct DDR pathways, with entirely different functions and enzymatic activities (see also Supplemental Table 1), suggesting that the loss of protein homeostasis that we observe is not limited to any particular molecular DDR defect. This is in line with other evidence linking defects in various DDR pathways to a loss of protein homeostasis (Arczewska et al., 2013; de Sousa Leal et al., 2020; Lee et al., 2018; Talaei et al., 2013).

The finding that the loss of protein homeostasis resulting from impairments in several DDR pathways is associated with an increase in the expression of HSP-16.2 mirrors our previous work, where we found an increase in the expression of HSPB5 in mammalian *ATM* KO cells (Huiting et al., 2021). Stable overexpression of HSPB5 in those cells was able to largely restore protein homeostasis, without affecting DNA repair capacity itself. This suggests that what we report here for *C. elegans* is analogous to our previous findings in human cells and indicates that HSPB5/αB-crystallin orthologs may represent a conserved second line of defense against the downstream proteomic consequences of loss of DDR capacity. Small heat shock proteins generally act as early interactors with stress-unfolded or misfolded proteins, keeping them in a state competent for either refolding by the HSP70/HSP90 machinery, or degradation by proteolytic systems (Mogk et al., 2019). HSPB5 has been reported to have a particularly broad substrate range, able to deal with many different destabilized protein species (Mymrikov et al., 2017). This suggests that impairments in certain DDR pathways may drive an increase in un- or misfolded proteins, the proteotoxic consequences of which can be buffered by elevated levels of HSP-16.2/HSPB5. Consistently, we find that in direct relation to safeguarding protein homeostasis, elevated HSP-16.2 expression is sufficient to prevent a motor impairment in *brc-1, atm-1* and *ung-1* compromised animals. These findings thus represent a proof of principle, as they connect the consequences of reduced DDR capacity to degeneration through a disruption of protein homeostasis. Although more work is needed to investigate the extent to which a loss of protein homeostasis indeed contributes to disease progression in DDR disorders in humans, our findings offer an attractive explanation for the degenerative phenotypes observed in these patients.

## MATERIALS AND METHODS

### Statistical analyses

All statistical tests were performed using Graphpad Prism software. In all protein insolubility and polyQ aggregation assays, a one-way ANOVA with Dunnet’s test was used to calculate significance compared to EV ctrl animals. For experiments with pairwise comparisons, Student’s unpaired t-test was used to calculate if two groups differed significantly, unless indicated otherwise. For behavioral experiments with multiple comparisons a one-way ANOVA with Dunnet’s test was performed to calculate significance compared to EV ctrl animals. All repetitions (n) originate from distinct samples (i.e. biological replicates).

### Media and strains

Unless mentioned otherwise, *C. elegans* were propagated at 20°C using standard lab conditions. Animals were age-synchronized by hypochlorite bleaching and eggs allowed to hatch overnight in M9 buffer. The strains used or generated in this study are listed in Table 1. In all experiments in Figures 2 and 3, a strain carrying a *rol-6* (HE1006) was used as a matched wild-type control for the *rol-6* integration marker in GL400 animals.

**Table 1.** Strains used in this study.

| Strain | Description | Genotype | Reference |
| --- | --- | --- | --- |
| N2 | Bristol wild isolate | Wild type |  |
| AM141 | Q40-YFP | rmls133[unc-54p::Q40::YFP] | (Morley et al., 2002) |
| HE1006 | Dominant Roller; matched wild type for GL400 | rol-6(su1006) II |  |
| GL400 | HSP-16.2::GFP | rfls400 [pCL79(hsp-16.2::GFP) + pRF4] | (McColl et al., 2010)<sup>†</sup> |

<sup>†</sup>A kind gift from dr. Gordon J. Lithgow

### RNAi experiments

Age-synchronized L1 larvae were cultured on NGM agar plates containing 1 mM IPTG and 50 mg/ml ampicillin seeded with RNAi bacteria until they reached adulthood. Adult animals were washed of the plates every other day and transferred to fresh RNAi plates to remove progeny. Washed animals were inspected visually to ensure no progeny was transferred. At adult day 5, animals were collected for further analysis via protein fractionation and RT-PCR. All RNAi constructs used in this study are from the Ahringer library, constructs are listed in Table 2. Knockdowns of DNA damage response (DDR) genes were included in the study only when *a priori* RT-PCR testing revealed a knockdown in N2 wild-type animals of >50% - only *msh-6, atm-1, rad-50, mre-11, brc-1* and *ung-1* passed this cut-off.

**Table 2.**
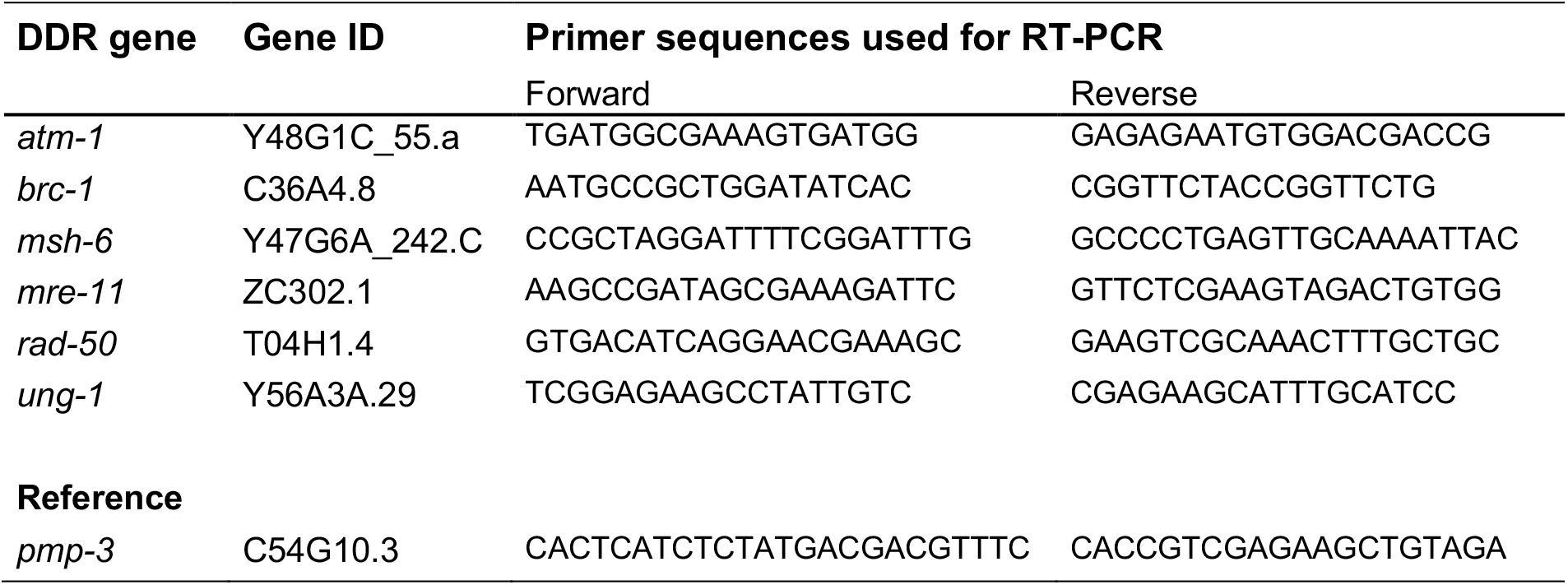
RNAi and primers.

### Quantitative RT-PCR in C. elegans

To assess the efficiency of RNAi knockdown in *C. elegans*, total RNA was extracted from RNAi treated animals using TRIzol reagent (#15596018, Invitrogen) according to the manufacturer’s instructions. The RNA concentration and quality was measured with a NanoDrop 2000 spectrophotometer (Thermofisher). cDNA was synthesized with the RevertAid H Minus First Strand cDNA Synthesis kit (#K1632, Life Technologies) using random hexamer primers. Quantitative real-time PCR was performed using a Roche LightCycler 480 Instrument II (Roche Diagnostics) with SYBR green dye (#172-5125, Bio-Rad) to detect cDNA amplification. Relative transcript levels were quantitated using a standard curve of pooled cDNA samples. Expression levels were normalized against *pmp-3;* an endogenous reference gene. Primers used for RT-PCR in this study are listed in Table 2.

### Protein fractionation

Animals were resuspended in lysis buffer (50 mM Tris/HCl pH 7.4, 100 mM NaCl, 1% v/v SDS, complete protease inhibitor cocktail (Roche Diagnostics)), homogenized by 5-7 20s cycles with a FastPrep-24 (MP Biomedicals) at 4 m/s, and clarified by low-speed centrifugation (2500 *rcf*). Protein content was measured and equalized (DC protein assay, Bio-Rad), Whole Cell Extract (WCE) fraction was taken and insoluble proteins were pelleted by high-speed centrifugation (21,000 *rcf*, 45 minutes). Insoluble protein pellets were washed with lysis buffer containing 50 mM Tris/HCl pH 7.4 and 100 mM NaCl, and solubilized in urea buffer (8 M urea, 2% v/v SDS, 50 mM DTT, 50 mM Tris/HCl pH 7.4) overnight at RT in a Thermomixer R (Eppendorf) at 1200 *rpm*. After extraction, insoluble, aggregated proteins were quantified by SDS-PAGE, in-gel silver staining with a Pierce™ Silver Stain Kit, and densitometry analysis using ImageJ software (FIJI).

### Western blotting

For Western blotting, proteins were transferred to a PVDF membrane, probed with anti-GFP (Takara Bio Clontech, #632380, mouse, 1:5000) and anti-Tubulin (Sigma, #T5138, mouse, 1:4000) as indicated, incubated with HRP-conjugated anti-mouse (GE Healthcare, #NXA931, 1:5000) and imaged in a Bio-Rad ChemiDoc imaging system.

### Quantification of polyglutamine aggregation

Polyglutamine aggregates present in Q40 animals fed RNAi bacteria were counted using a fluorescence dissection stereomicroscope (Leica Microsystems). Twenty animals were counted per experiment and each experiment was repeated for at least 3 times. Punctae were counted with no size cut-off.

### Germline apoptosis

Synchronized young adult animals (AD2-3) grown on RNAi bacteria at 25° were stained with 75 µg/ml acridine orange (AO)(Sigma) for 1 hour, and transferred to fresh NGM agar plates for 3 hours to clear excess AO from their intestines. AO positive germ corpses present in the posterior gonads were counted using a fluorescence dissection stereomicroscope (Leica Microsystems), for each indicated condition 40 animals were counted. Irradiated animals were analyzed 24-36 hours post irradiation with the indicated doses (IBL-637 irradiator, CIS Biointernational).

### Larval development assay

Adult day 2 RNAi-fed animals reared at 25°C were allowed to lay eggs on corresponding fresh RNAi NGM agar plates for 2 hours. After 50 hours, the developmental stage of offspring was evaluated based on vulval morphology and the presence of eggs to distinguish young adults from gravid adult animals.

### Crawling assay

Adult day 5 animals RNAi-fed bacteria reared at 25°C were transferred onto fresh 9 cm NGM agar plates, 8 animals per plate and two plates per RNAi, using fixed marked starting positions. Animals were allowed to crawl overnight. The next day, animals were taken off, and plates were imaged and subsequently analyzed using a digital grid overlay to quantify crawling tracks.

### Paralysis assay

Each individual paralysis experiment was started with 100 L4 RNAi-fed animals and carried out at 25°C. Animals were placed on fresh RNAi plates (10 worms per plate) and were assessed daily for paralysis by touching their noses and tail-prodding with a paintbrush hair. Worms that moved their nose but not their body were scored as paralyzed.

## ACKNOWLEDGEMENTS

We thank Gordon Lithgow for kindly providing us with the GL400 strain. This work was supported by a Nederlandse organisatie voor Wetenschappelijk Onderzoek (NWO) grant to SB [ALW 824.15.004], and by generous funding from Charity4Brains.

## AUTHOR CONTRIBUTIONS

W.H. and S.B. conceived and designed the study. E.A.A.N. and H.H.K. contributed to scientific hypotheses generation. W.H. and A.D.J. performed protein fractionations. R.I.S. performed RT-PCR experiments. W.H. performed all other experimental work. W.H. and S.B. interpreted the data. W.H. generated figures. W.H. and S.B. wrote the manuscript. All authors critically reviewed the manuscript.

## FIGURES AND LEGENDS

**Supplemental Figure 1.**
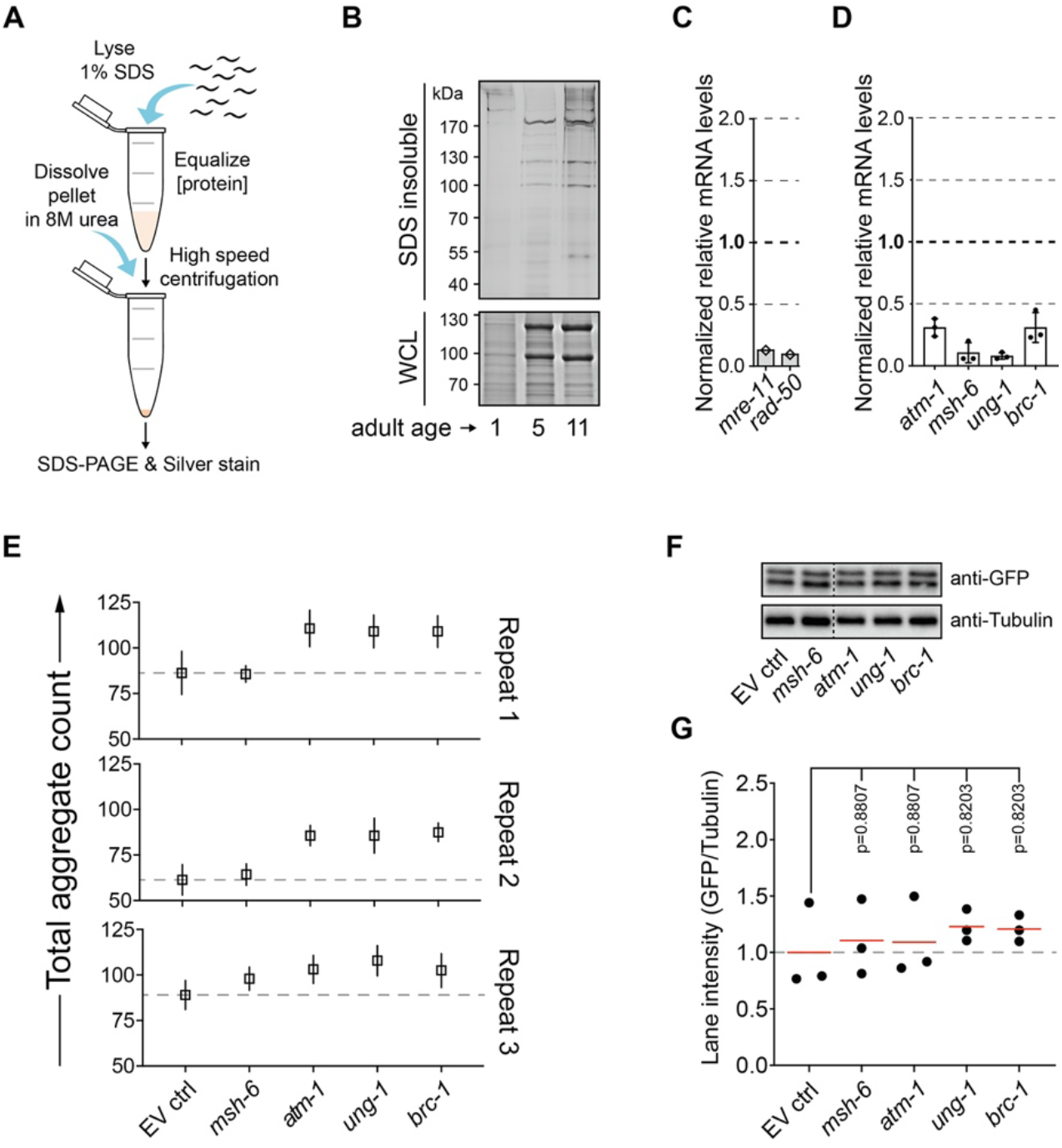
Protein aggregation is accelerated in *C. elegans* after targeting of various DDR components. (A) Overview protein fractionation procedure. (B) SDS-insoluble and WCL protein fractions of animals of the indicated age, run on SDS-PAGE gels and subjected to in-gel silver or Coomassie staining, respectively. (C, D) Knockdown efficiencies of the indicated genes measured with RT-PCR. (E) Absolute Q40::YFP aggregate counts of three independent repeats. (F) Western blot analysis of AM141 (Q40::YFP) animals subjected to the empty vector (EV ctrl) or indicated RNAi’s, probed with anti-GFP and anti-Tubulin. Dotted line indicates a cropping. (G) Densitometric quantification of three independent repeats of F. Significance calculated using a one-way ANOVA with Dunnet’s post-hoc test.

**Supplemental Figure 2.**
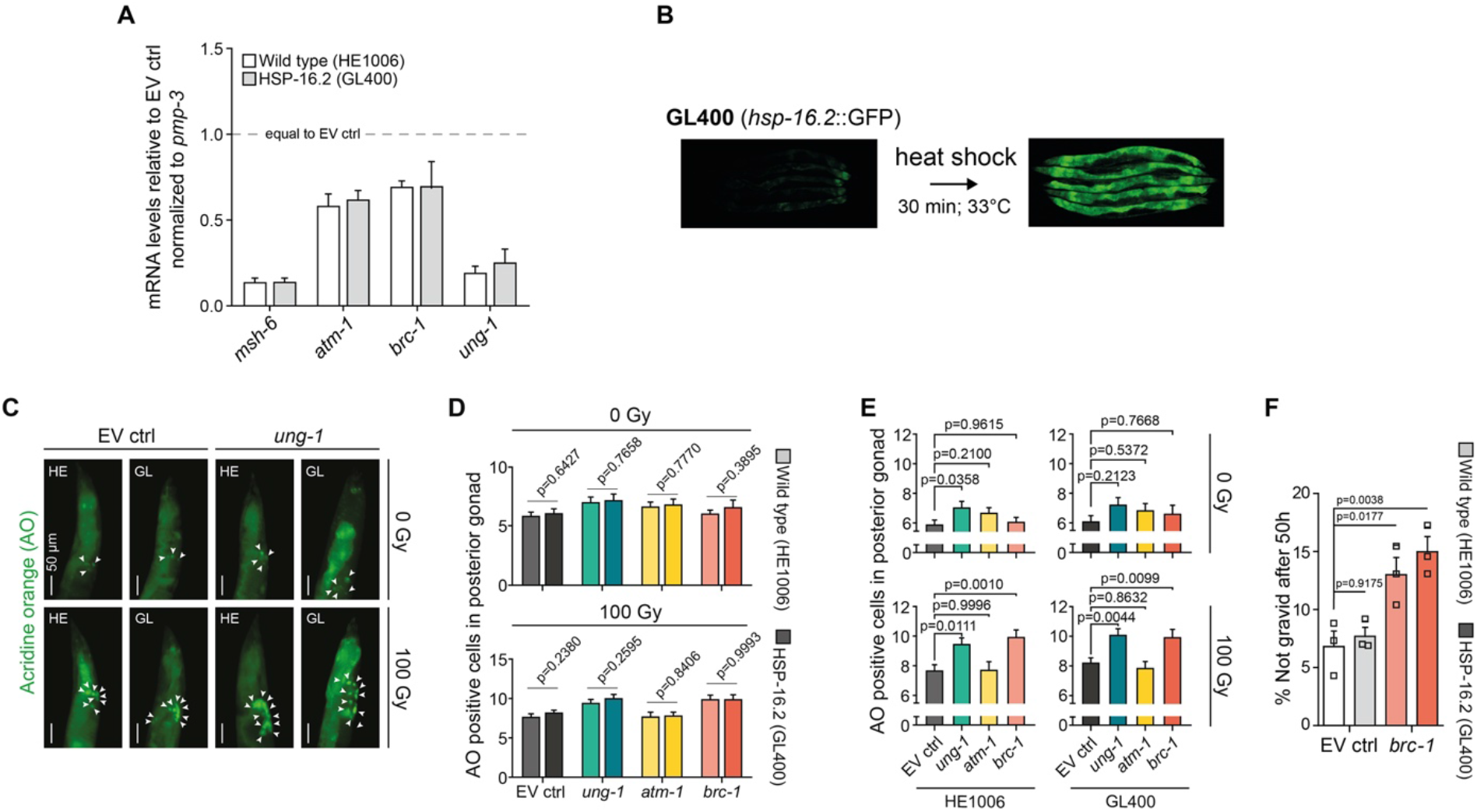
HSP-16.2 overexpression does not affect genome maintenance capacity. (A) RT-PCR of RNAi-fed wild-type (HE1006) and HSP-16.2 overexpression (GL400) animals, relative to EV ctrl. Data represented as mean ± SEM. (B) HSP-16.2::GFP signal before and after heat shock in GL400 animals. Typical fluorescence stereomicroscopy pictures are shown. (C) Fluorescence stereomicroscopy pictures of the posterior gonads in indicated animals, γ-irradiated with 100 Gy or not, stained with acridine orange. Arrow heads indicate acridine orange positive apoptotic bodies. Typical fluorescence stereomicroscopy pictures are shown. (D) Graphs depicting the quantification of C; wild-type against HSP-16.2 animals. Significance calculated using Student’s t-tests. (E) Graphs depicting the quantification of C; EV ctrl against the indicated RNAi’s, with or without gamma irradiation. Significance calculated using one-way ANOVAs with Dunnet’s post-hoc test. (F) Graph depicting progeny delay in wild-type and HSP-16.2 animals, fed empty vector (EV ctrl) or *brc-1* RNAi bacteria, 50h after synchronization. Y-axis shows percentage of animals not yet developed into gravid adults. Significance calculated using one-way ANOVAs with Dunnet’s post-hoc test. In all graphs, data is represented as mean ± SEM.

**Supplemental Figure 3.**
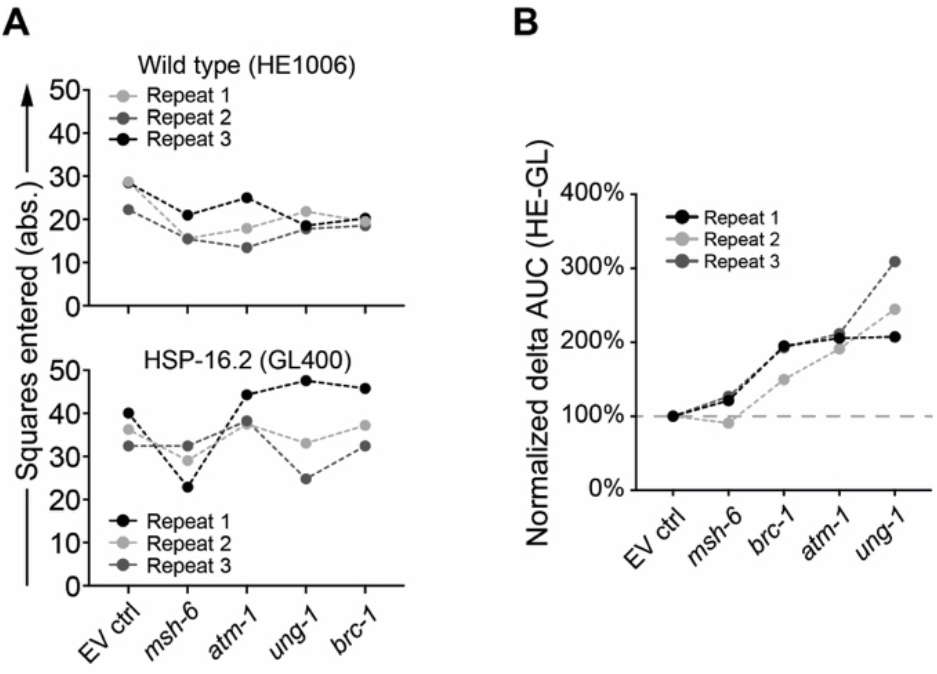
HSP-16.2 prevents a health span decline after knockdown of *atm-1, ung-1* or *brc-1*. (A) Absolute quantification of crawling tracks of HE1006 (wild-type) and GL400 (HSP-16.2 overexpression) animals fed the indicated RNAi’s, see also Figure 3. (B) Difference in area under the curve (AUC) between paralyis rates of HE1006 (wild-type) and GL400 (HSP-16.2 overexpression) animals fed the indicated RNAi’s.

